# A Paper-based Loop-Mediated Isothermal Amplification (LAMP) Assay for Highly Pathogenic Avian Influenza

**DOI:** 10.1101/2024.08.12.607641

**Authors:** Mohamed Kamel, Josiah Levi Davidson, Mohit S. Verma

## Abstract

Avian influenza outbreaks have had significant economic and public health consequences worldwide. Therefore, prompt, reliable, and cost-effective diagnostic devices are crucial for scrutinizing and confining highly pathogenic avian influenza viruses (HPAIVs). Our study introduced and evaluated a novel paper-based loop-mediated isothermal amplification (LAMP) test for diagnosing the H5 subtype of the avian influenza virus (AIV). We meticulously designed and screened LAMP primers targeting the H5-haemagglutinin (H5-HA) gene of AIV and fine-tuned the paper-based detection assay for best performance. The paper-based LAMP assay demonstrated a detection limit of 500 copies per reaction (25 copies/µL). This inexpensive, user-friendly point-of-need diagnostic tool holds great promise, especially in resource-limited settings. It only requires a water bath for incubation and enables visual detection of results without special equipment. Overall, the paper-based LAMP assay provides a promising method for rapidly and reliably detecting the H5 subtype of AIV, contributing to improved surveillance and early intervention strategies.

## 1 Introduction

Highly pathogenic avian influenza (HPAI) outbreaks in recent years have led to significant financial losses for the global poultry industry and pose a significant threat to public health. Furthermore, in recent years, HPAI has been found to infect a variety of animals other than avian populations, including seals, polar bears, goats, dairy cows, and humans, and as of Fall 2023, has now been detected in populations on every continent except Australia ^1^. An outbreak of H5N1 HPAI in Indiana led to over $100 million in poultry losses and triggered trade restrictions in 2023 ^2^. Similarly, the 2022 avian influenza outbreak in the US resulted in the loss of 40 million animals and cost $2.5 to $3 billion ^3^. Globally, avian influenza outbreaks in 2022 affected 67 countries, resulting in the loss of 131 million domestic poultry. Recent data reveals that avian influenza has also impacted dairy cattle, goats, and other livestock, with a notable multi-state outbreak of H5N1 HPAI reported in dairy cows for the first time in March 2024 ^4–6^. These outbreaks underscore the far-reaching economic and public health risks posed by avian influenza viruses (AIVs) in poultry and beyond, affecting various sectors of agriculture and livestock ^7,8^.

Influenza viruses belong to the Orthomyxoviridae family and have four types (A, B, C, and D) with unique antigenic properties. Influenza A viruses are further classified into subtypes based on their surface glycoproteins: hemagglutinin (HA) with 18 subtypes and neuraminidase (NA) with 11 subtypes, from which 16 HA subtypes and 9 NA were identified in birds ^9–16^.

AIVs, specifically influenza A, are commonly found among birds and can be categorized as high or low pathogenicity (HPAIV and LPAIV) ^17–19^. H5 and H7 AIV subtypes have the potential to mutate to HPAIV in galliform avian species, leading to high mortality and outbreaks ^20^.

Various diagnostic tests, including polymerase chain reaction (PCR)-based techniques like reverse transcription PCR (RT-PCR) and reverse transcription quantitative PCR (RT-qPCR), are used to identify AIVs ^21–26^. However, these traditional and molecular methods have limitations, such as being time-consuming and requiring sophisticated facilities ^27^. There is an urgent need for a rapid and efficient diagnostic approach for AIV infection ^28–31^. Loop-mediated isothermal amplification (LAMP) is one such alternative that provides a precise and sensitive approach for RNA/DNA amplification at a constant temperature ^32,33^. LAMP is resistant to inhibitors, cost-effective, and suitable for point-of-need testing. By utilizing LAMP assays, we can detect and track the prevalence of H5 subtype AIV and achieve early detection of infected patients.

The LAMP reaction only requires a water bath to be conducted in non-laboratory settings. It can detect RNA templates by using reverse transcriptase and DNA polymerase ^32^. Reverse transcription (RT)-LAMP techniques have been successfully developed to detect various RNA viruses, including the severe acute respiratory syndrome coronavirus 2 ^34^, West Nile ^35^, dengue ^36^, and others ^37–39^. For AIV, Hu *et al*. (2017) utilized the HA gene from H5 influenza A viruses in digital LAMP (dLAMP) using microfluidic droplet technology and successfully detected 10 copies/μL of the target *in vitro* transcribed (IVT) sequence ^40^. Table 1 summarizes information on the development of LAMP assays targeting different subtypes of AIV. The AIV target genes, LOD, template type, and detection methods used in previous LAMP assays were compiled and presented. Detection methods included real-time fluorescent assays, turbidity assays, gel electrophoresis, digital LAMP, visual observation by colorimetry, integration with lateral flow assays (LFA), and microarrays.

**Table 1:**
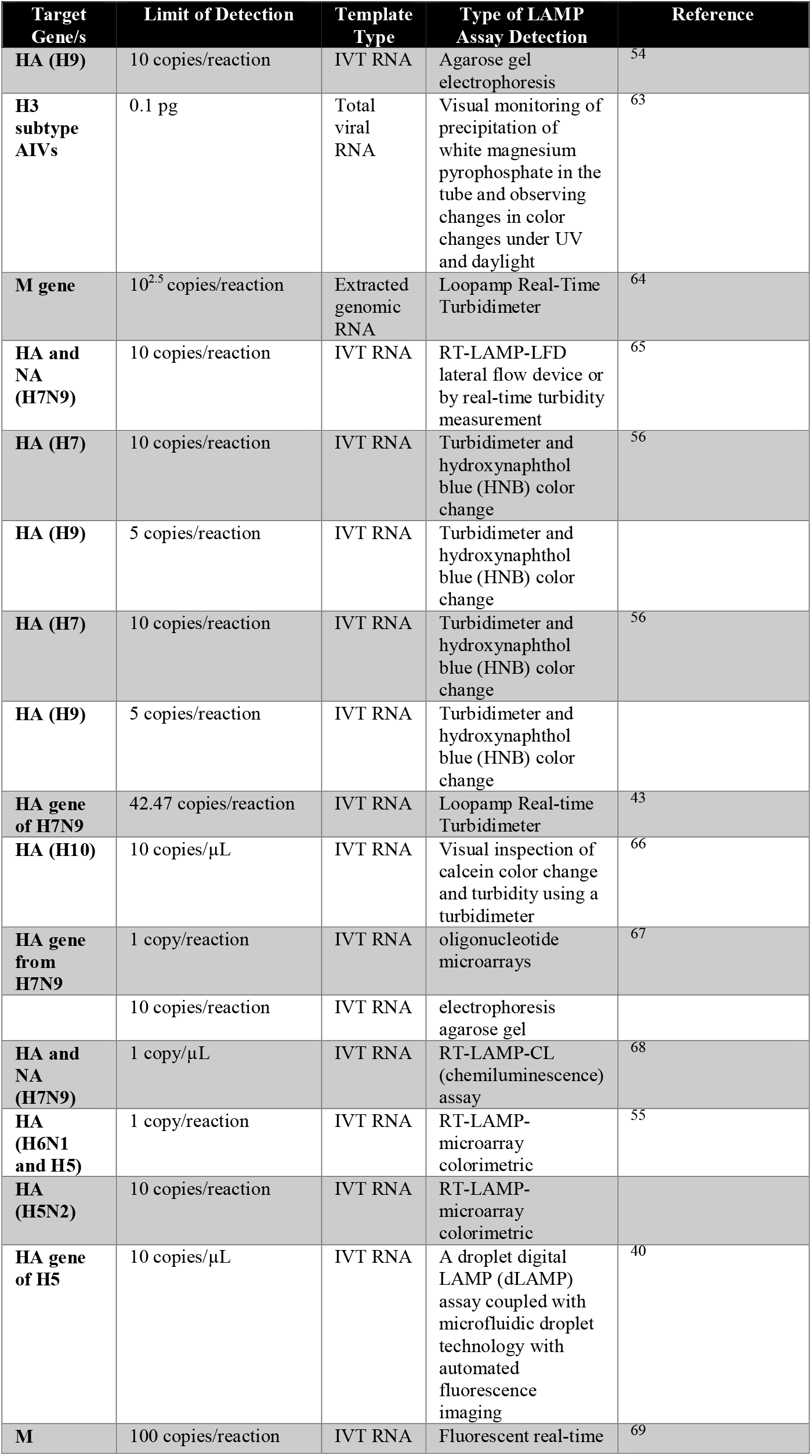
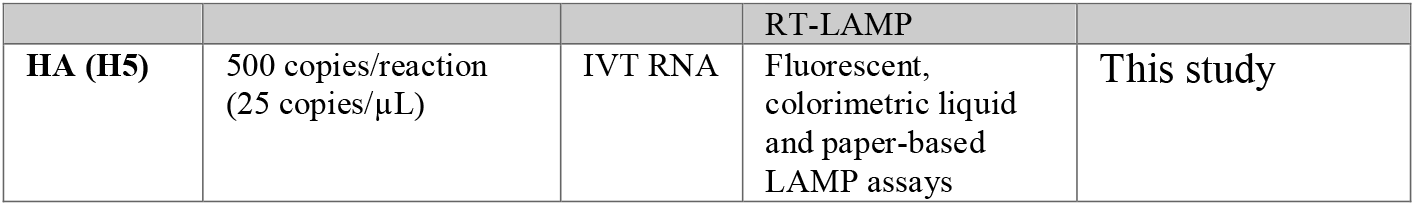
The LAMP assays developed so far targeting different AIVs and their subtypes have been presented in the Table, showing their target genes, LOD, type of template, and the used detection method. The selection criteria for including LAMP assay studies here in the table were based on the reported viral load quantification in copies, picograms, or nanograms. This criterion was chosen to maintain consistency and comparability across studies.

In our study, the RT-LAMP system implemented on a paper-based device can detect H5 subtype AIV at a minimum of 500 copies per reaction. These microfluidic paper-based analytical devices (µPADs) are user-friendly, portable, and customizable, making them ideal for rapid AIV infection diagnosis in both laboratory and field settings. By employing user-friendly and portable colorimetric liquid and paper LAMP assays, we can use these assays as screening tools for rapid AIV infection diagnosis. This approach has the potential to significantly enhance influenza A virus detection and contribute to the control and prevention of avian influenza outbreaks.

## 2 Results

### 2.1 LAMP primer screening

The present study reports the results of LAMP primers designed to target the HA segment of an Influenza A H5N1 virus isolated from Turkey in Indiana, United States in 2022 using a sequence made available by GISAID ^41^. Furthermore, a multiple alignment of various H5 subtype AIV HA sequences from NCBI was used to obtain conserved sequences for primer design. The design process involved using Primer Explorer v5 (http://primerexplorer.jp/e) and the addition of degenerate primers. Primer sets were designated as HPAIV.HA.*x* where HPAI refers to the target organism, HA refers to the target gene, and *x* is an arbitrary index that increases by one for each new primer set designed (Table 2). Primer screening was conducted using fluorometric LAMP with a double-stranded DNA intercalating dye that is spectrally similar to SYTO9 (**Error! Reference source not found**.). We used four technical replicates using 10^5^ copies per reaction of IVT synthetic RNA of the HA gene in the positive reactions and nuclease-free water in the no-template control (NTC) reactions to conduct the screening. Only primer sets without amplification in the NTC (indicating the presence of false positives) were chosen for further analysis. Notably, we observed that primers HA.11 and HA.12 were unsuitable for further analysis, as they yielded false positives. On the other hand, the primer sets HPAIV.HA.1 through HPAIV.HA.10 displayed no false positives, along with higher fluorescence intensity and successfully amplified positive controls. The most effective LAMP assay primer sets were determined by subjecting the various sets to identical conditions, including primer concentration (final concentration of 1.6 µM FIP/BIP, 0.2 µM F3/B3, 0.4 µM LoopF/B), reaction mixture (25 μL), and reaction program (65 °C), using the same template input (10^5^ copies/reaction). The performance of these primers was assessed by measuring maximal fluorescence intensity, rapid response time, and the absence of any false positives within negative controls. Of the proficient primers, the H5 subtype HPAIV.HA.4, H5 subtype HPAIV.HA.5, H5 subtype HPAIV.HA.6, and H5 subtype HPAIV.HA.7 sets demonstrated the highest level of amplification efficiency, as evidenced by scoring results (Table 3).

**Table 2:**
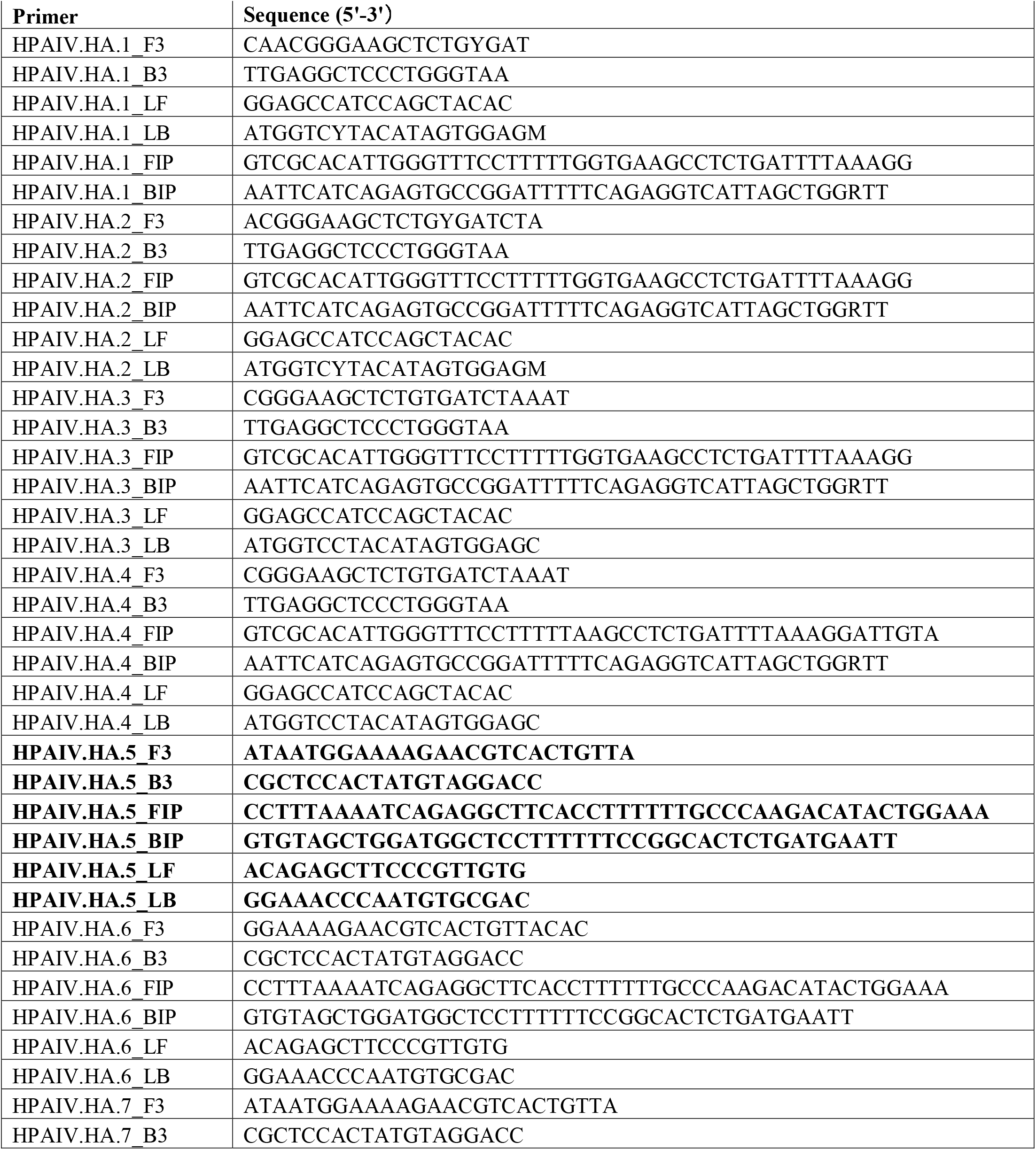
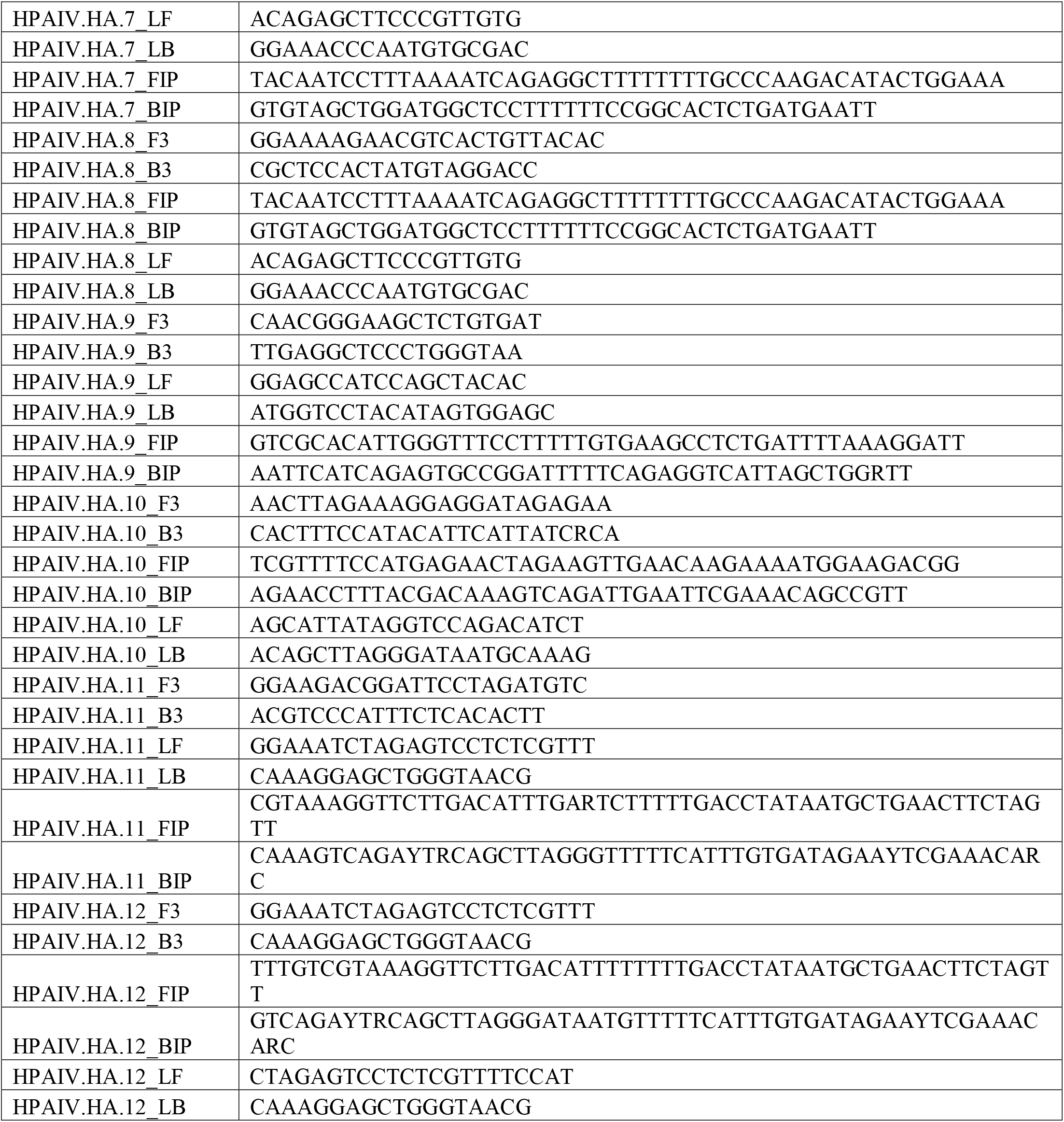
List of RT-LAMP primers designed and used in this study for the HA target gene.

**Table 3:**
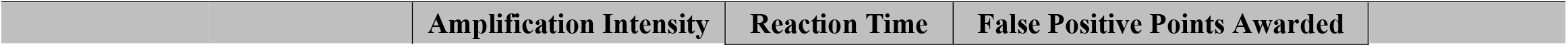
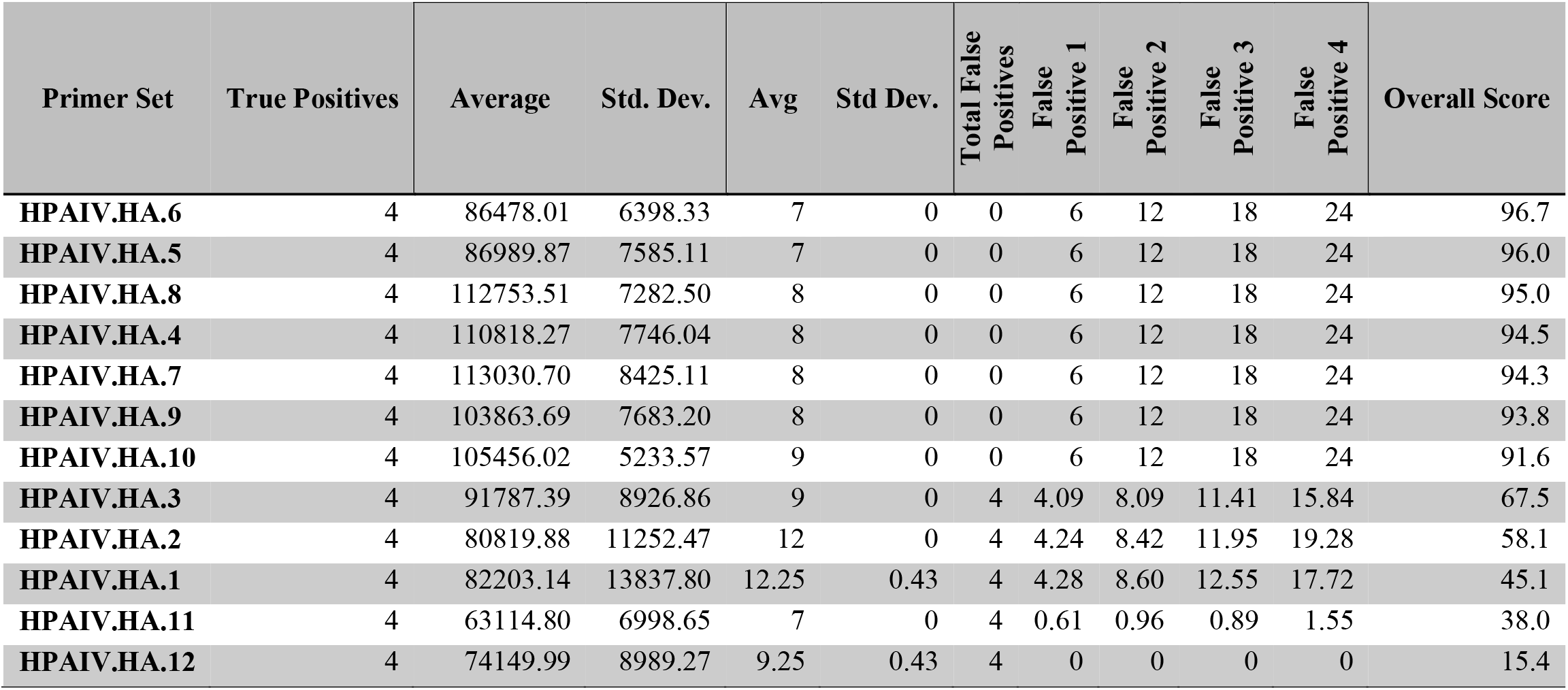
Performance characteristics of designed LAMP primer sets for screening purposes to select the best-performing LAMP primer sets for further downstream assays.

|  | Amplification Intensity | Reaction Time | False Positive Points Awarded |
| --- | --- | --- | --- |

| Primer Set | True Positives | Average | Std. Dev. | Avg | Std Dev. | Total False<br>Positives | False<br>Positive 1 | False<br>Positive 2 | False<br>Positive 3 | False<br>Positive 4 | Overall Score |
| --- | --- | --- | --- | --- | --- | --- | --- | --- | --- | --- | --- |
| HPAIV.HA.6 | 4 | 86478.01 | 6398.33 | 7 | 0 | 0 | 6 | 12 | 18 | 24 | 96.7 |
| HPAIV.HA.5 | 4 | 86989.87 | 7585.11 | 7 | 0 | 0 | 6 | 12 | 18 | 24 | 96.0 |
| HPAIV.HA.8 | 4 | 112753.51 | 7282.50 | 8 | 0 | 0 | 6 | 12 | 18 | 24 | 95.0 |
| HPAIV.HA.4 | 4 | 110818.27 | 7746.04 | 8 | 0 | 0 | 6 | 12 | 18 | 24 | 94.5 |
| HPAIV.HA.7 | 4 | 113030.70 | 8425.11 | 8 | 0 | 0 | 6 | 12 | 18 | 24 | 94.3 |
| HPAIV.HA.9 | 4 | 103863.69 | 7683.20 | 8 | 0 | 0 | 6 | 12 | 18 | 24 | 93.8 |
| HPAIV.HA.10 | 4 | 105456.02 | 5233.57 | 9 | 0 | 0 | 6 | 12 | 18 | 24 | 91.6 |
| HPAIV.HA.3 | 4 | 91787.39 | 8926.86 | 9 | 0 | 4 | 4.09 | 8.09 | 11.41 | 15.84 | 67.5 |
| HPAIV.HA.2 | 4 | 80819.88 | 11252.47 | 12 | 0 | 4 | 4.24 | 8.42 | 11.95 | 19.28 | 58.1 |
| HPAIV.HA.1 | 4 | 82203.14 | 13837.80 | 12.25 | 0.43 | 4 | 4.28 | 8.60 | 12.55 | 17.72 | 45.1 |
| HPAIV.HA.11 | 4 | 63114.80 | 6998.65 | 7 | 0 | 4 | 0.61 | 0.96 | 0.89 | 1.55 | 38.0 |
| HPAIV.HA.12 | 4 | 74149.99 | 8989.27 | 9.25 | 0.43 | 4 | 0 | 0 | 0 | 0 | 15.4 |

### 2.2 Fluorescent LAMP LOD

The LOD of the RT-LAMP assay for the detection of H5 subtype AIV was evaluated by testing four selected primer sets (H5 subtype HPAIV.HA.4, H5 subtype HPAIV.HA.5, H5 subtype HPAIV.HA.6, and H5 subtype HPAIV.HA.7) for the fluorescent LOD study. The qLAMP method was also tested by diluting the IVT template to evaluate its ability to detect the lowest number of copies. The LODs of these primer sets ranged from 250 to 18900 copies per reaction. The observed LOD for the primer sets H5 subtype HPAIV.HA.4, 5, 6, and 8 were 500, 500, 18900, and 1890 copies/reaction, as shown in Fig. 2, Supplementary Fig.**Error! Reference source not found**., Supplementary Fig. **Error! Reference source not found**., and **Error!**

**Fig. 1:**
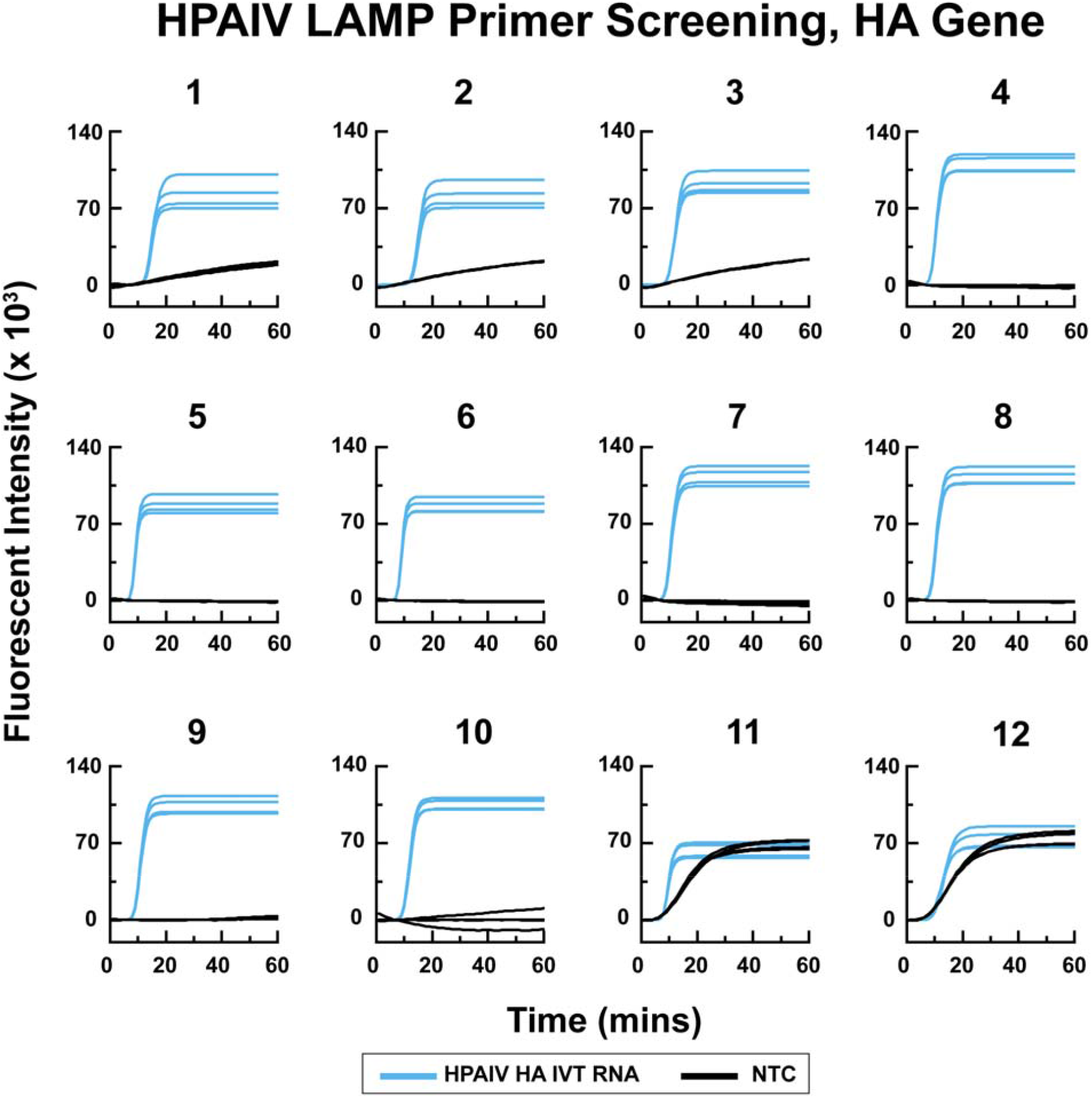
Screening of LAMP primer sets for H5 subtype AIV by fluorescent LAMP assay using the NEB Warm Start DNA/RNA LAMP Kit.

**Fig. 2:**
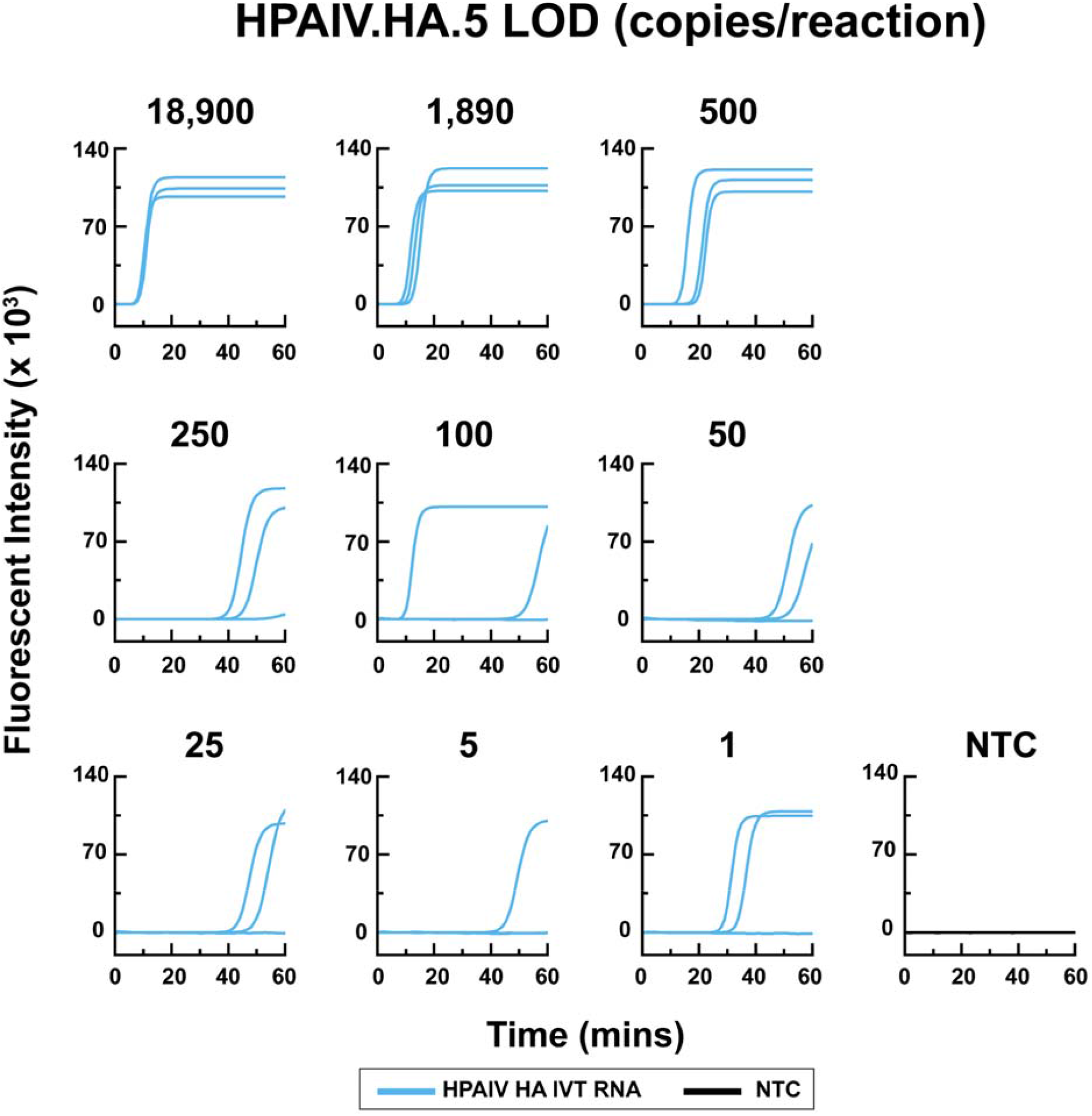
LOD experiment showing the result of the H5 subtype AIV qLAMP assay of the HPAIV.HA.5 primer set using the fluorescent qLAMP assay testing different concentrations of IVT RNA (18900, 1890, 500, 250, 125, 50, 25, 5, and 1 copies/reaction)

**Reference source not found**.. The assay exhibited a LOD of 500 copies/reaction, indicating its potential for effective detection of H5 subtype AIV.

### 2.3 Colorimetric liquid LAMP LOD

The two primer sets (HPAIV.HA.4 and HPAIV.HA.5) that exhibited a better fluorescent LOD compared to the other evaluated primer sets were further evaluated using a colorimetric liquid LOD method. Both primer sets’ colorimetric assay prompted a pink to yellow color change. The H5 subtype HPAIV.HA.5 primer set showed an LOD of 500 copies/reaction, while the other primer set (H5 subtype HPAIV.HA.4) exhibited a higher LOD than 1890 copies/reaction (Fig. 3 and **Error! Reference source not found**.). The results indicate that the H5 subtype HPAIV.HA.5 primer set is the preferred primer set for H5 subtype AIV detection.

**Fig. 3:**
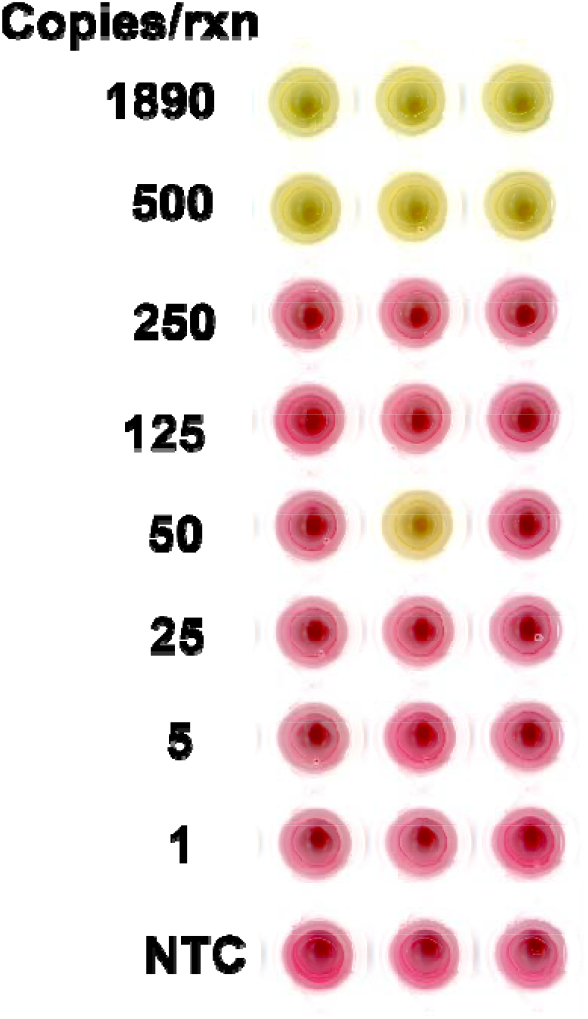
Limit of Detection (LOD) experiment of the H5 subtype AIV colorimetric liquid LAMP assay using the HPAIV.HA.5 primer set. The colorimetric liquid LAMP assay was performed to test different concentrations of synthetic RNA (1890, 500, 250, 125, 50, 25, 5, and 1 copies/reaction). The figure shows a LOD of 500 copies/reaction for the H5 subtype AIV colorimetric liquid LAMP assay using the HPAIV.HA.5 primer set, where the color changes from pink to yellow.

### 2.4 Paper-based LAMP LOD

The study also assessed the efficiency of the paper-based LAMP assay using the same primer set H5 subtype HPAIV.HA.5 at varying template concentrations. The paper-based LAMP assay’s LOD was comparable to that of the colorimetric liquid LAMP assay, indicating its reliable detection ability with a low detection limit (500 copies/reaction; Fig. Fig. 4).

**Fig. 4:**
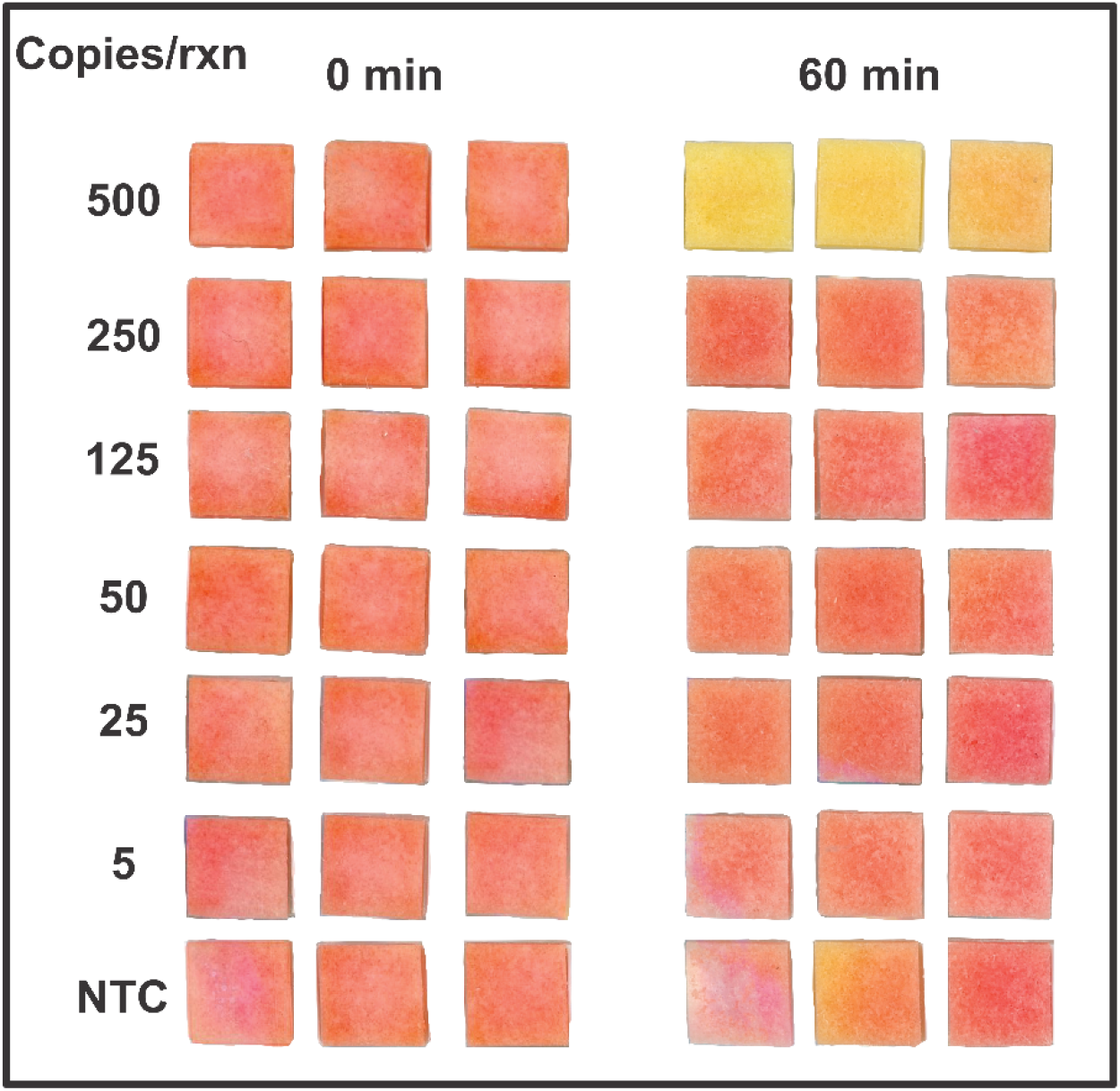
Paper-based LAMP LOD: Limit of Detection (LOD) experiment of the H5 subtype AIV colorimetric paper-based LAMP assay using the HPAIV.HA.5 primer set. The colorimetric paper-based LAMP assay was performed to test different concentrations of synthetic RNA.

The results of LAMP primer screening and detection limit testing demonstrate the optimized LAMP assay’s potential for detecting H5 subtype AIV. The study’s findings have significant implications for clinical diagnosis and epidemiological studies of H5 subtype AIV. The optimized LAMP assay can prove to be an essential diagnostic tool for the effective control of H5 subtype AIV through early detection.

## 3 Discussion

AIVs present a significant threat to the global poultry industry, causing substantial mortality and economic damage. These viruses can infect various bird species, raising concerns about animal-to-human transmission and potential fatalities ^42–44^. The recent HPAI H5N1 outbreak in U.S. dairy cattle highlights the virus’s ability to jump from wild birds to domestic animals and humans, underscoring the risk of a future influenza pandemic ^45–47^. Developing sensitive screening methods, such as LAMP alongside traditional techniques, is crucial for containing the spread of the virus and safeguarding poultry populations ^48^. Point-of-need tests (PONT) offer promise for on-site sample testing without requiring transportation of the samples to the lab, with LAMP being particularly suitable for remote areas with limited lab facilities. This study focuses on developing efficient and expeditious assays utilizing PONT RT-LAMP assays that detect AIV manifest with approximately ten times the detection capability of the endorsed RT-PCR method.

Our RT-LAMP assay exhibits a LOD of 500 RNA copies per reaction (25 copies/µL), which is better than conventional RT-PCR assays with LOD ranging from 1000 to 10,000 copies per reaction ^49,50^ but worse than recent RT-qPCR assays that report LOD of 10 copies per reaction ^51^. Furthermore, our study also surpasses multiplex RT-qPCR assays like that of Le *et al* which had a detection limit of 50 copies per microliter, equivalent to 1,000 copies per reaction ^52^ and the matrix gene RT-qPCR assay by Spackman *et al*. which has an LOD of 10 fg, approximately 125 copies per microliter of the target RNA ^53^. While acknowledging the superior LOD achieved by certain real-time RT-qPCR assays ^51^, it is crucial to note that these methods may not be suitable for field deployment. Our RT-LAMP assay strikes a balance between limit of detection and practicality, making it a valuable tool for the rapid identification of AIV.

The development of RT-LAMP primer sets targeting the H5-HA gene was a primary focus of this study. To assess the efficacy of these primers, *in vitro* transcribed RNA fragments of the H5-HA gene were used for experimental validation. The rationale behind adopting this methodology was driven by the unfeasibility of obtaining field-isolated AIV samples due to regulatory constraints surrounding the pathogen as it was a select agent. As such, we developed our RT-LAMP primers using synthetic RNA of a conserved region found in AIVs obtained from varied poultry populations around the world. This finding indicates that our RT-LAMP assays would be effective in detecting H5-HA presence.

Several studies have evaluated the performance of the LAMP assay for AIV detection ^54– 56^. The detection of AIVs has been explored through the use of the LAMP method, resulting in promising sensitivity for detection, with a specific focus on the H5N1 and H7 subtypes. A RT-LAMP method developed for the detection of H7N9 subtype AIV showed high specificity and sensitivity, with an LOD of up to 50 copies/reaction and could be used without the need for extra nucleic acid extraction steps using a direct RT-LAMP ^43^.

Our assays show promise for surveillance screening and detection of the H5 subtype AIV in clinical samples, including cloacal and throat swabs, as previously reported for the H9 subtype ^54^. The H9N2 LPAIV load in various cloacal swab and oropharyngeal samples varied from 4.1 × 10 to 7.7 × 10^5^ RNA copies per microliter, with the highest viral loads in the cloacal swab and oropharyngeal samples being 1.0 × 10^5^ and 7.7 × 10^5^ RNA copies per microliter, respectively ^57^. Together, these studies demonstrate the potentially high utility of our assay in detecting a clinically relevant amount of HPAIV in field-based settings.

Paper-based LAMP assays are a promising diagnostic tool for detecting AIVs. One of their main advantages is their simplicity and ease of use, as they do not require sophisticated laboratory infrastructure or trained personnel. This makes them suitable for resource-limited settings ^58,59^. Moreover, these assays are highly specific and sensitive, as demonstrated by the development of a highly robust and effective RT-LAMP assay for detecting H5 subtype AIV. This assay showed a LOD of 500 copies/reaction, indicating its high diagnostic potential for AIVs’ detection.

In addition to their current diagnostic capabilities, paper-based LAMP assays have promising prospects. One such prospect is the development of multiplex LAMP assays, which can simultaneously detect multiple target analytes. This includes the multiplex RT-LAMP assay, which offers quick, highly sensitive, inexpensive, and robust diagnostics for detecting multiple AIVs. Another potential future application of paper-based LAMP assays is their integration with digital technologies, like smartphone apps, for real-time monitoring and analysis. This integration could lead to the creation of affordable PONTs usable in both clinical and non-clinical settings, such as field surveillance of AIV in wild bird populations.

While the RT-LAMP assay offers numerous advantages, it is important to acknowledge its drawbacks as well. The constant genetic variability of AIV and the potential for non-specific amplification can lead to inaccurate positive results. Moreover, the assay’s performance (in terms of speed, limit of detection, and specificity) may be compromised by the complex sample matrix and the presence of inhibitors. However, these challenges can be overcome by regularly updating the primer sequences to detect new outbreaks and using highly specific primers to minimize false positives. Furthermore, assay performance can be enhanced by employing straightforward techniques like heat or chemical treatment, as well as sample dilution.

## 4 Conclusions

We developed a platform capable of detecting AIV using fluorescent and colorimetric liquid and paper LAMP assays that produce colorimetric responses visible to the naked eye. Our platform has the following seven advantages: i) it demonstrates an LOD of 500 copies/reaction (25 copies/μL), ii) it requires minimal operator training, iii) it is simple to use, iv) it requires only a water bath for operation, v) it provides a colorimetric response visible to the naked eye, vi) it is suitable for resource-limited settings, and vii) it is the first paper-based LAMP assay for detecting HPAI.

Four limitations and their opportunities for enhancing the current strategy are: i) LAMP-based assays are prone to false positives due to aerosol contamination and nucleic acid product generation, which can be minimized by implementing advanced sealing procedures, ii) an external heater is still needed for conducting the assay, but future versions could feature an integrated battery-operated heater to enhance portability and convenience, iii) individual variations in color perception could affect the interpretation of results, which can be addressed by using smartphones or cameras for precise color quantification, and iv) sample distribution across reaction zones can be uneven, but developing a spreading layer could ensure even sample distribution.

Due to the simplicity and scalability of this test, we envision that it could be used in a wide variety of settings, including in-field diagnostics. Our platform can be readily adapted for multiplexing different HA genes to detect a wider range of influenza types and subtypes, as well as other pathogens. The reconfigurable nature of our platform makes it an ideal tool for the deployment and detection of emerging pathogens in future public health emergencies.

To fully utilize these potential advantages, additional research is necessary. This includes validating our findings with clinical samples and assessing the feasibility of implementing these assays in real-world scenarios.

## 5 Methods

### 5.1 Plasmids and in vitro transcription

The HA segment of AIV type A/H5N1 (GISAID Accession #EPI1985974) of viral isolate A/turkey/Indiana/22-003707-003/2022 (|HA|4|WSS3056019) was used as a template for the synthetic gene sequence. The H5 subtype HPAIV.HA was synthesized and cloned into the pBluescript II KS (+) plasmid provided by Genescript Biotech under the stream of the T7 promoter sequence. The sequence had been terminated by the T7 terminator. SacI and NotI restriction endonuclease recognition sequences were placed upstream and downstream of the HA gene, respectively. The recombinant plasmid carrying the target HA gene insert was then linearized by the restriction enzyme NotI-HF^®^ (NEB R3189; Ipswich, MA, USA) using rCutSmart buffer for 15 min. The linearized plasmid DNA template was then subjected to a plasmid purification step using the Wizard Clean-up System (Promega A7280; Madison, WI, USA). The purified linearized DNA, including the T7 promoter, was then used as a template for *in vitro* transcription using HiScribe^®^ T7 Quick High Yield RNA Synthesis Kit (NEB E2050; Ipswich, MA, USA) per the supplier’s instructions. The *in vitro* transcribed RNA was subjected to 2 µl of DNAase enzyme treatment for 30 min in DNase buffer to eliminate any remnants of the template DNA. The *in vitro* transcribed RNA was purified using Monarch RNA Clean-up Kits (NEB T2030; Ipswich, MA, USA). Subsequently, the nanodrop 8000 spectrophotometer (Thermo Scientific, Wilmington, Delaware) was used to measure the RNA concentration.

### 5.2 Primer design and synthesis

LAMP primer sets were designed and prioritized over the conserved portions of the HA sequence of HPAI virus type A/H5N1 (GISAID Accession #EPI1985974) utilizing the LAMP Primer Explorer v5 program (http://primerexplorer.jp/e; Eiken Chemical) or manual methods following the instructions of Primer Explorer v5 program. The sequences were subjected to multiple sequence alignment using Clustal W by default setting, and some primers were modified as degenerate primers manually (Table 2). The feasibility of all primer sets was then validated using the BLAST program (BLASTN). All designed primer sets included loop primers (LF/LB) to increase reaction speed. The LAMP primer sets for the HA gene shown in Table 2 were synthesized and desalted by Invitrogen (Waltham, MA, USA).

### 5.3 Fluorescent LAMP assays

The fluorescent LAMP assay was carried out with the primers listed in Table 2. The master mix utilized for the assay was composed of 2.5 µl of 10× HA LAMP primer mix, 12.5 µl of WarmStart® 2X Master Mix E1700 LAMP Kit (NEB, USA), and 5 µl 5× LAMP fluorescent dye (diluted in nuclease-free water from 50X LAMP fluorescent dye provided with the kit) per reaction. Subsequently, 20 µl of this mixture was dispensed evenly into a 96-well plate (Thermo Scientific, AB-0800/W), followed by adding 5 µl of purified RNA (at varying concentrations) or molecular grade nuclease-free water in the case of NTC reactions. The plate was then capped using Versicap mat cap stripts (Fisher Scientific, Catalog No AB1820100). The sealed plate was then placed in a qTower 3G Touch (Analytic Jena, Germany) or qTower 3G () and incubated at 65 °C with a ramp rate of 0.1 °C/sec for 60 minutes. Real-time fluorescence was detected on the blue channel using settings for the FAM dye. Fluorescence measurements were taken every 60 seconds during the reactions.

### 5.4 LAMP screening and scoring

Primer sets were screened by performing 4 replicates of positive control reactions containing 5 µL of 2.0 × 10^3^ copies/reaction of synthetic HA gene IVT RNA and varying the primer set. In the case of NTC reactions, 5 µL of nuclease-free water was used in place of synthetic HA gene IVT RNA. LAMP reactions were carried out as detailed in the entitled “Fluorescent LAMP assays”.

Primer sets were scored according to a weighted schema previously designed by our lab ^60^. Briefly, primer sets were scored based upon the average reaction time, standard deviation of reaction time, average maximum fluorescent intensity, and standard deviation of maximum fluorescent intensity across all 4 replicates, as well as the number of false positives with weights of 20%, 15%, 5%, 5%, and 55% of the primer set score, respectively (Table 3). The reaction time for an individual reaction was defined as the time at which the second derivative of the time rate of change of the fluorescent intensity reached a maximum (i.e., roughly when the exponential phase is first beginning).

False positives were determined as any signal in an NTC reaction that displayed non-negligible fluorescent intensity increases over the course of the reaction and that increased above 10% of the maximum intensity of the instrument (approximately 140,000 RFU) in a negative reaction. False positives were penalized with increasing severity as the number of false positives increased. Primer sets were then sorted based on the total score. Primer sets scoring above 95 were selected and one additional primer set was selected in the event that downstream screening failed. Information on accessing the code for primer scoring can be found in the section entitled “Data “.

### 5.5 Limit of detection experiment for LAMP assay

The LODs for candidate primer sets were established using serial dilutions of HA-IVT in water based on its dilutions of 1980, 500, 250, 125, 50, 25, 5, and 1 copies/reaction in 25 μL reactions. A benchtop (Epson Perfection V800 Photo Color) scanner was used to scan the plate to capture the colors of the reaction mixtures at different time points in the colorimetric liquid LAMP. An assessment of LOD is based on a primer set with the lowest virus concentrations, resulting in a distinct color change across all three replicates in colorimetric liquid/paper-based LAMP or exhibiting a fluorescent amplification curve in the three triplicates.

### 5.6 Colorimetric liquid LOD assay

New England BioLabs® Inc. has developed a pH-dependent master mix that incorporates the highly modified BST 2.0 WarmStart DNA Polymerase in a special buffer containing essential co-factors MgSO4 and phenol red as a visible pH-sensitive indicator. This indicator changes from pink to yellow when the pH of the reaction mixture drops. This is due to the large amount of proton produced by BST 2.0 WarmStart DNA polymerase due to extensive nucleic acid amplification activity. LAMP results were more visible and detectable with this type of master mix. DNA polymerase I from Bacillus stearothermophilus has been genetically engineered to be highly active in an isothermal environment without the need for a denaturation step. The WarmStart Master Mix is designated because the enzyme is inactive at lower temperatures but becomes active once the reaction solution reaches a temperature of 40°C (NEB, USA). Using the enzyme and indicator of this recent invention, a simple device can be used for incubation, and the color of the amplicon could be detectable by the naked eye within a short time. Since the reaction was performed in a water bath at 65°C, all samples were incubated simultaneously for detection without needing a thermocycler. In addition, color detection of the amplicon simplified the reading of the results, eliminating the need for a post-amplification processing step.

The colorimetric assay was performed according to the instructions in the NEB WarmStart Colorimetric LAMP 2X Master Mix Kit containing UDG (NEB M1804). This mix provides a distinctive red-to-yellow transition in the presence of a pH indicator dye to indicate a positive result. Reactions consisted of 2.5 μL 10× primer mix (1.6 μM FIP and BIP primers, 0. 4 μM each of LF and LB primers, 0.2 μM each of F3 and B3 primers), 12.5 μL WarmStart Colorimetric LAMP 2X Master Mix, 5 μL nuclease-free water, and 5 μL template RNA or molecular biology water for NTCs in a total volume of 25 μL. All reactions were then incubated in a qPCR cycler at 65°C for 60 minutes with the lid heated to 95°C. During the RT-LAMP procedure, the Bst 2.0 WarmStart® DNA Polymerase (NEB, USA) was fortified with dUTP to avert carryover contamination. Furthermore, the antarctic thermolabile UDG (NEB, USA) was employed to eliminate DNA contamination and degrade any dU-containing DNA. The WarmStart Colorimetric LAMP 2X Master Mix (DNA and RNA) (NEB, USA) is comprised of a specialized reaction mixture incorporating a modified and WarmStart strand-shifting DNA polymerase (Bst 2.0) and reverse transcriptase (RTx), enables rapid and highly reliable detection of both DNA (LAMP) and RNA (RT-LAMP).

### 5.7 Colorimetric paper-based LAMP assay

We used a formulation previously described by our group for the on-paper colorimetric assay to detect SARS-CoV-2 ^58,61^: Final colorimetric paper-based RT-LAMP master mix included MgSO_4_ (8 mM), KCl (50 mM), WarmStart® RTx reverse transcriptase (0.3 U/μL), *Bst* 2.0 WarmStart® DNA polymerase (0.32 U/μL), antarctic thermolabile UDG (0.0004 U/μL), dUTP (0.14 mM), dNTP mixture (1.4 mM each dNTP), phenol red (0.25 mM), tween® 20 (1% v/v), trehalose (10% w/v), BSA (500 μg/mL), and betaine (20 mM). Upon device assembly, 25 µL of the aforementioned master mix was added to each pad along with the appropriate primer set as specified in the section entitled “Colorimetric liquid LOD assay”. The devices were then left for 1 hour to dry, after which 20 µL of either RNA-free water or a solution containing RNA was added to each pad individually for rehydration. The rehydrated devices were then placed in 2 × 2 inch 2 mil polypropylene bags (ULINE S-17954). To maintain a constant temperature of 149°F (or 65°C), a 12-quart EVERIE Sous Vide Container (Amazon B07GQWP85C) was filled with water and an Anova Culinary AN500-US00 Sous Vide Precision Cooker (Amazon B07WQ4M5TS) was used. Each appliance bag was securely attached to a standard clear film (617993, Office Depot, USA) and then placed inside a larger 1-gallon Savour Sous Vide Cooking Bag (Amazon B07NCSXMNN). After heating the water bath, the sealed and capped cooking bag was inserted. The samples were allowed to remain in the water for 60 minutes. Then, images for the experiments were captured at 0 and 60 minutes with an Epson Perfection V800 Photo Scanner (Amazon B11B223201) configured to Pro Mode, 48-bit color image type, with a resolution of 600 dpi.

### 5.8 Fabrication and optimization of devices for paper-based LAMP assay

This device comprises reading and reaction layers and spacers that prevent crosstalk among strips. The 3 mil optically clean MELINEX (Tekra MELINEX[r] 454 polyester [PET]) backing was used to create the reading area. The double-sided adhesive was adhered to the reaction strips, which were 5 mm by 5 mm of chromatography paper (Ahlstrom-Munksjö Grade 222). These strips were then separated by the 2.5 x6 mm 20-mil polystyrene spacers (Tekra Double White Opaque HIPS Polystyrene Litho Grade). 20 µl of water was added to saturate the strips during rehydration.

## 7 Acknowledgements

We gratefully acknowledge all data contributors, i.e., the authors and their originating laboratories responsible for obtaining the specimens and their submitting laboratories for generating the genetic sequence and metadata and sharing via the GISAID Initiative, on which this research is based.

This work was supported by the Foundation for Food and Agriculture Research under award number – Grant ID: FF-NIA20-0000000087. The content of this publication is solely the responsibility of the authors and does not necessarily represent the official views of the Foundation for Food and Agriculture Research. Josiah Levi Davidson was also supported by Grant Number, UL1TR002529 (S. Moe and S. Wiehe, co-PIs) from the National Institutes of Health, National Center for Advancing Translational Sciences, Clinical and Translational Sciences Award.

## 8 Author contributions

M.K. performed the experiments, prepared the original manuscript draft, prepared the figures, and analyzed the data. J.L.D. ran the scoring code and prepared the figures. M.S.V obtained funding and supervised the work. MK, J.L.D, and M.S.V reviewed and edited the manuscript drafts. All authors approved the final version of the manuscript.

## 9 Data availability statement

Raw data collected in preparation of this manuscript are available via Mendeley Data at https://www.doi.org/10.17632/34xgdvvv3m.1 ^62^.

The code used during primer scoring can be found online at the Purdue University Github at https://github.itap.purdue.edu/VermaLab/PrimerScoring. Additionally, a release of the code is included in the raw data archive via Mendeley data above.

## 10 Competing interests

M.S.V. has interests in Krishi Inc., which is a startup company developing molecular assays. Krishi Inc. did not fund this work. Remaining authors do not have a competing interest.

## 11 Declaration of AI and AI-assisted technologies in the writing process

During the preparation of this work, the authors used Grammarly (https://grammarly.com/) to check for grammar errors and improve the academic writing language. After using this tool/service, the authors reviewed and edited the content as needed and take full responsibility for the content of the publication.

