## Supporting Information for "A Paper-based Loop-Mediated Isothermal Amplification (LAMP) Assay for Highly Pathogenic Avian Influenza"

**for**

**Supplementary figures**

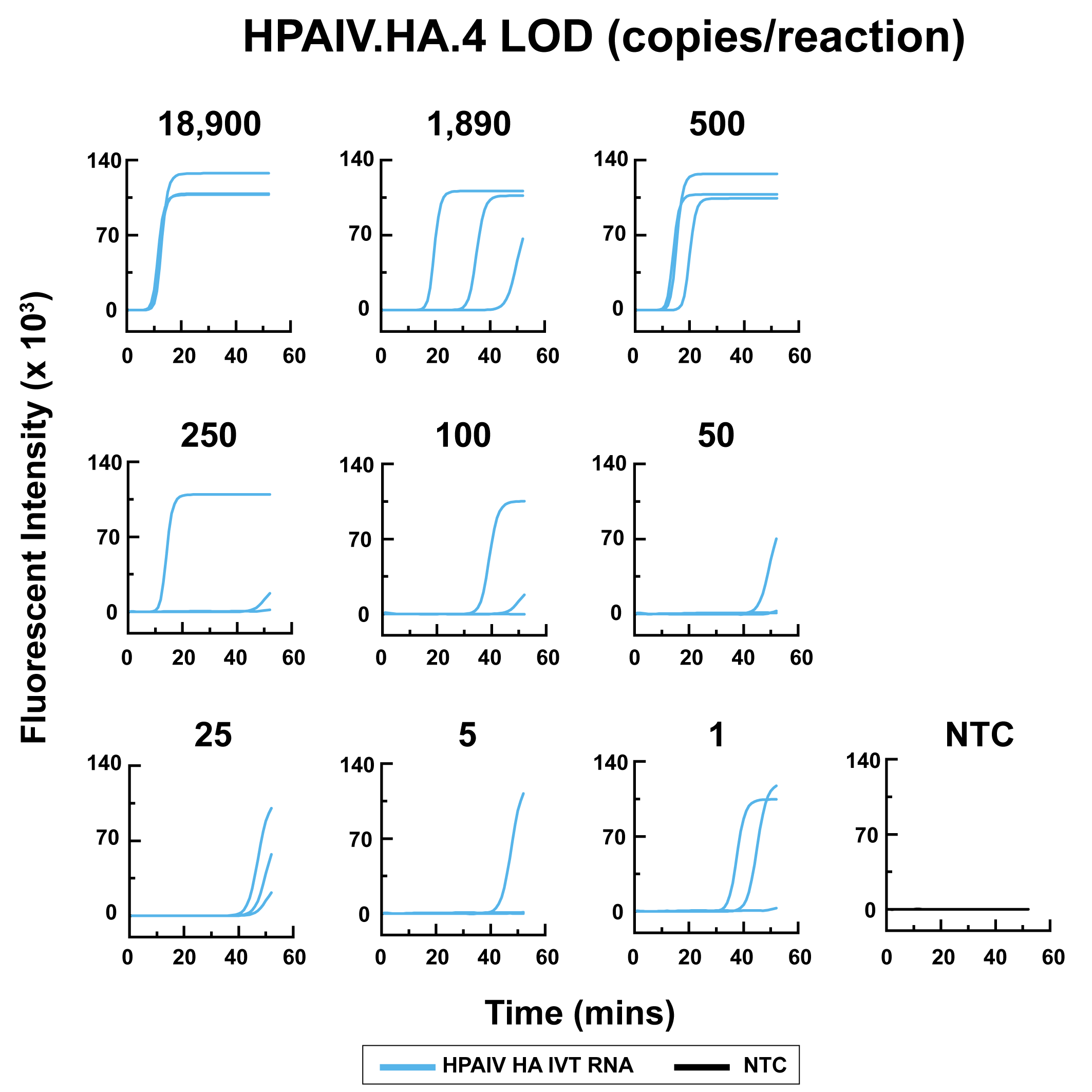

Supplementary Fig. 1: LOD experiment showing the result of the H5 subtype AIV qLAMP assay of the HPAIV.HA.4 primer set using the fluorescent qLAMP assay testing different concentrations of IVT RNA (18900, 1890, 500, 250, 125, 50, 25, 5, and 1 copies/reaction)

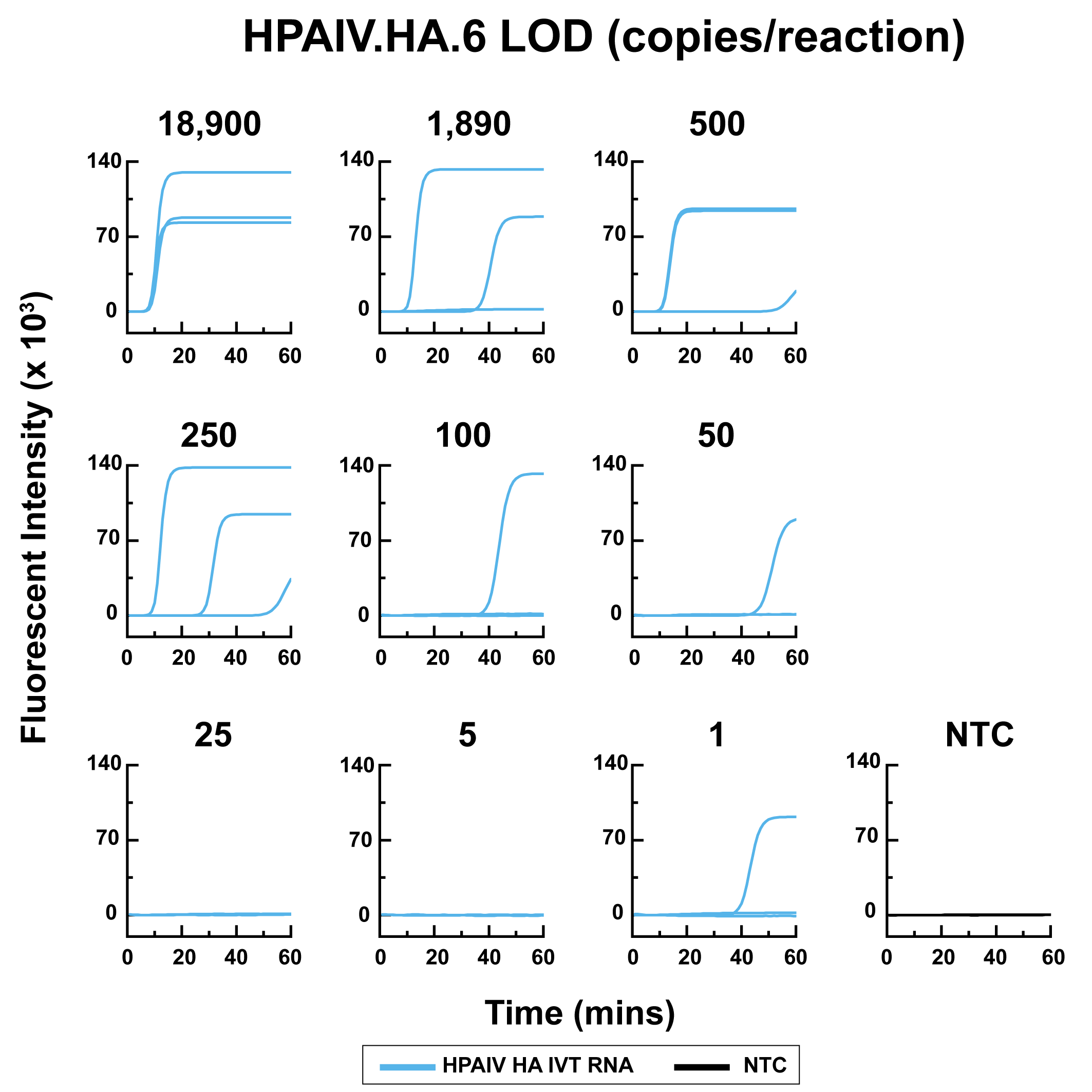

Supplementary Fig. 2: LOD experiment showing the result of the H5 subtype AIV qLAMP assay of the HPAIV.HA.6 primer set using the fluorescent qLAMP assay testing different concentrations of IVT RNA (18900, 1890, 500, 250, 125, 50, 25, 5, and 1 copies/reaction)

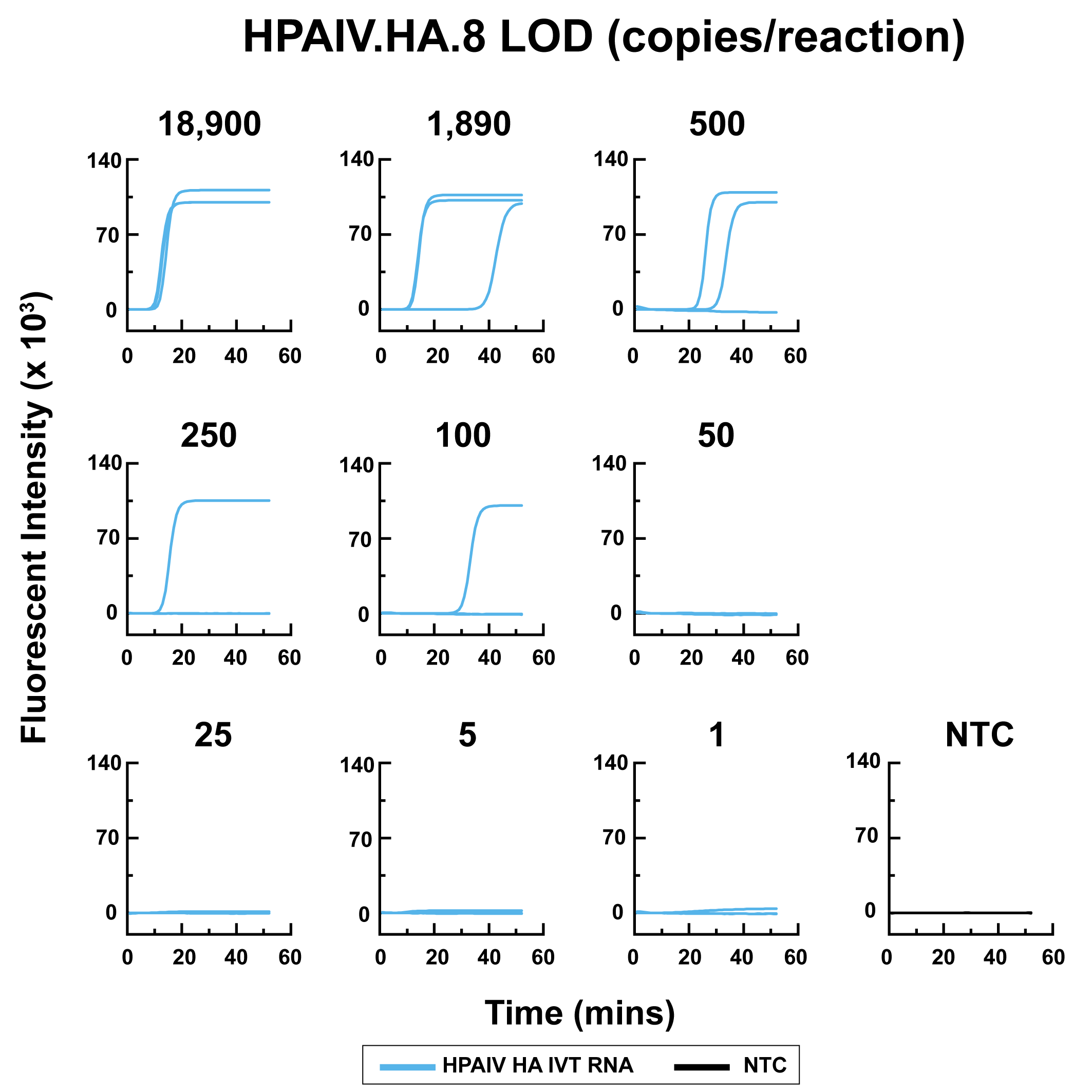

Supplementary Fig. 3: LOD experiment showing the result of the H5 subtype AIV qLAMP assay of the HPAIV.HA.8 primer set using the fluorescent qLAMP assay testing different concentrations of IVT RNA (18900, 1890, 500, 250, 125, 50, 25, 5, and 1 copies/reaction)

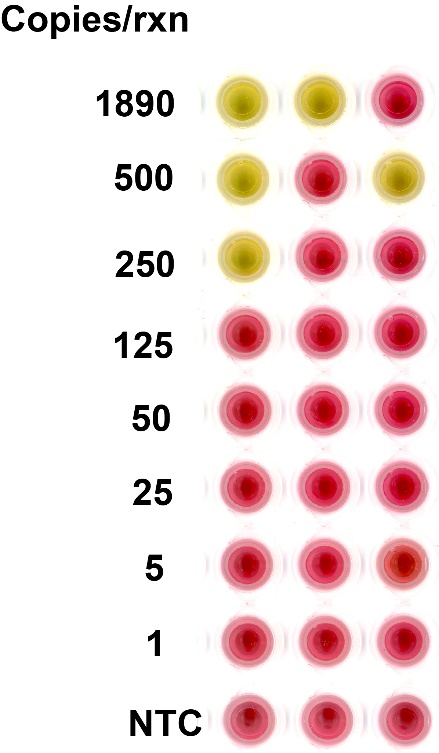

Supplementary Fig. 4: Limit of Detection (LOD) experiment of the H5 subtype AIV colorimetric liquid LAMP assay using the HPAIV.HA.4 primer set. The colorimetric liquid LAMP assay was performed to test various concentrations of IVT RNA (1890, 500, 250, 125, 50, 25, 5, and 1 copies/reaction).
